# Sexual selection does not predict long-term population trends in birds

**DOI:** 10.64898/2026.05.18.725879

**Authors:** Miguel Gómez-Llano, Chris Cooney, Tim Janicke, Robert MacDonald, Edward H. Morrow

## Abstract

Sexual selection is a major evolutionary force, yet its demographic consequences remain unclear. While experimental studies often report positive effects of sexual selection on traits linked to population performance, comparative studies often find null or negative associations with population persistence. One explanation for this discrepancy is that the demographic consequences of sexual selection depend on ecological context, particularly variation in mortality and fecundity. Here, we used six decades of abundance data and test whether sexual selection predicts population trends across 738 bird species from Europe and North America. We quantify sexual selection using complementary proxies capturing different components of sexual selection: mating system, sexual dichromatism, sexual size dimorphism and relative testes mass. We further assess whether the effect of sexual selection in population trends is mediated by mortality and fecundity. Across all proxies, we found no evidence that sexual selection is associated with population trends. This result is consistent across continents and robust to variation in mortality and fecundity. Our findings suggest that, despite its central role in shaping phenotypic evolution, sexual selection does not translate into consistent effects on long-term population trends at macroecological scales. More broadly, these results highlight a potential disconnect between evolutionary processes and population dynamics.

## Introduction

Sexual selection is a major evolutionary force shaping phenotypes, behaviour, and reproductive strategies across taxa^1^. Sexual selection can promote the spread of beneficial alleles accelerating adaptation^2–6^ and protecting against extinction^7^. However, it may also impose demographic costs through traits that reduce survival^1,8,9^ or intensify sexual conflict^10–12^. As a result, the net consequences of sexual selection for long-term population trends remain unresolved. Determining whether sexual selection enhances or undermines population viability is therefore key to linking evolutionary processes with large-scale patterns of biodiversity and population change.

Experimental studies have approached this question by comparing populations under different mating systems (e.g., monogamy versus polyandry). A meta-analysis of laboratory studies reported a small but positive effect of sexual selection on traits associated with population fitness^13^. In contrast, studies of wild populations frequently find weak, null, or negative associations between sexual selection and extinction risk, persistence, or population trends^14–19^. This discrepancy may arise because laboratory studies typically reduce environmental heterogeneity and constrain natural selection, conditions under which the benefits of sexual selection may be more readily expressed^20–23^. In addition, experimental systems are often biased toward short-lived, highly fecund organisms^13^, in which evolutionary responses may be more easily detectable over relatively short generations. This bias could make the beneficial effects of sexual selection easier to observe. Differences in how sexual selection is quantified may also contribute: comparative studies often rely on proxies such as sexual size dimorphism or dichromatism, which may capture only specific components of sexual selection^24,25^ or be confounded with natural selection^26^. Using multiple complementary proxies, including those more directly comparable to experimental manipulations, may therefore help reconcile results across approaches.

A key but less frequently tested explanation is that the demographic consequences of sexual selection depend on ecological context. In particular, mortality and fecundity are expected to influence the opportunity for selection and thus whether the effects of sexual selection can be expressed at the population level^20,23,27^. Elevated and stochastic mortality, especially during early life stages, can reduce variance in reproductive success and the opportunity for selection, potentially offsetting any benefits of sexual selection^27,28^. In contrast, higher fecundity may buffer populations against the demographic costs of sexual selection, facilitating adaptation and the spread of beneficial alleles^20,23^. These contrasting effects suggest that the relationship between sexual selection and long-term population trends may be context dependent and depend on how they are mediated by survival and fecundity. Yet, empirical tests of how demographic factors modulate this relationship remain scarce.

Here, we analyzed long-term population trends for 738 bird species from Europe and North America to test whether sexual selection predicts long-term population trends. We used long-term population trends as they are one of the best predictors of extinction risk^29^. We quantify sexual selection using four complementary proxies capturing different components of pre- and postcopulatory selection: mating system (a proxy similar to treatments used in laboratory experiments), sexual dichromatism (precopulatory), sexual size dimorphism (precopulatory), and relative testes mass (postcopulatory). In addition to testing overall associations between sexual selection and long-term population trends, we evaluate two theory-driven hypotheses. First, we test whether elevated mortality reduces the demographic consequences of sexual selection. We used migratory status (resident and migratory) as a proxy for mortality because (1) mortality increases during migration and is higher in offspring^30^, and (2) in partial migratory populations (with resident and migratory individuals) residents have a survival advantage over migratory^31^. Thus, we expected that resident species with less mortality would show a more positive effect of sexual selection on population trends than migratory species. Second, we test whether higher fecundity enhances the beneficial effects of sexual selection^23^ by examining interactions with clutch size and clutch number. By integrating multiple proxies within a single phylogenetically controlled comparative framework, we assess whether the relationship between sexual selection and long-term population trends depends on the type of sexual selection and on the demographic context in which it operates.

## Methods

### Population trends

Birds are among the best-studied animal groups and provide a unique opportunity to analyze long-term population trends at large spatial scales. Population trends for European species were obtained from the Pan-European Common Bird Monitoring Scheme, which provides species-specific trend estimates (slope and standard error) from the 1980s to 2024^32^.

For North American species, population trends were estimated from count data from the North American Breeding Bird Survey, a standardized long-term monitoring program with annual counts from the 1960s to 2023^33^. To ensure comparability with the European data, we reconstructed species-specific trends using site-level time series. We defined sites as unique combinations of country, state, and route, analogous to monitoring sites in PECBMS. We used the full set of surveyed route-year combinations (“weather.csv”) and, for routes in which a species was recorded at least once, treated years with no detections as zero counts. For each species, we fitted a log-linear Poisson model with the number of individuals as the response variable and site and year (centered on the initial survey year for each species) as fixed effects. We then extracted the exponential of the year coefficient as an estimate of the long-term population trend, along with its standard error, to match the format of the European data. Thus, values of population trends of one represent stable populations, below and above show declining and increasing populations, respectively. In total, the dataset includes 738 species (171 from Europe and 567 from North America), with 15 species present in both datasets.

### Sexual selection

We quantified sexual selection using four complementary proxies capturing different components of pre- and postcopulatory selection: mating system, sexual dichromatism, sexual size dimorphism and relative testes mass. Mating system (monogamy versus polygamy) was based on the classification by Barber et al.^34^, whom scored species on a scale from 0-4 as 0 = strict monogamy, 1 = frequent monogamy, 2 = regular polygamy, 3 = frequent polygamy, and 4 = extreme polygamy. We converted this into a binary index as 0 = monogamy and 1 = polygamy (encompassing 1-4 in^34^) to provide a measure comparable to experimental manipulations of mating system^7^.

Sexual dichromatism was used as a measure of pre-copulatory sexual selection. To quantify the degree of sexual dichromatism within species, we followed a previous study^35^ and used the summed Euclidean distances between sex-specific measurements of plumage colouration extracted from a large dataset of calibrated images of museum specimens (http://www.projectplumage.org/). Briefly, plumage patch locations on specimen images were first located using a trained deep convolutional neural network model^36^ and then used to extract calibrated reflectance measurements for 10 discrete body regions across up to 3 replicate male and female specimens for each species. Reflectance measurements were averaged across replicates and then used to calculate cone catch (*u, s, m, l, dbl*) values based on a typical avian ultraviolet-sensitive visual model^35,37^. Finally, chromatic cone catch values (*u, s, m, l*) were mapped into tetrahedral colour space using functions available in *pavo 2*^38^ and then combined with *dbl* cone measurements to calculated Euclidean distances between male and female patch measurements. A dichromatism score of zero indicates identical colouration in both sexes (monochromatism) with higher positive values indicating greater degree of dichromatism.

We quantified sexual size dimorphism as (mass of the larger sex / mass of the smaller sex) − 1^39^, using body mass data from published databases^40–42^. Although sexual size dimorphism may reflect both natural and sexual selection, it is widely used as a comparative proxy in macroecological studies^25,43^. Correlations among proxies were low (all r < 0.22; Table 1), indicating that they capture largely independent aspects of sexual selection. Finally, we used residual testes mass^34^ as a proxy for post-copulatory sexual selection. Residual testes mass captures variation in investment in sperm competition and is weakly correlated with mating system.

**Table 1.** Phylogenetically controlled correlations between the different proxies of sexual selection. Because data of these proxies were available for different set of species, we did pairwise phylogenetically controlled correlations using phylogenetic generalized least squares on the standardised values of each proxy. Shown are correlation coefficients (*r*) and number of species in brackets.

|  | Residual testes<br>mass | Sexual<br>dichromatism | Sexual size<br>dimorphism |
| --- | --- | --- | --- |
| Mating system | 0.217 (316) | 0.012 (566) | 0.114 (563) |
| Residual testes mass |  | 0.082 (324) | 0.083 (332) |
| Sexual dichromatism |  |  | 0.002 (578) |

### Mortality and fecundity

To test whether mortality modulates the effects of sexual selection, we used migratory status (migratory or resident^34^) as a proxy for differences in survival. Migratory species experience elevated and often stochastic mortality, particularly during migration and early life stages^30^, which may reduce the opportunity for selection. To test whether fecundity influences the effects of sexual selection, we used clutch size (number of eggs per clutch) and clutch number (number of clutches per year), obtained from Myhrvold et al.^44^.

### Analysis

We tested whether sexual selection predicts long-term population trends using multilevel meta-analytic models implemented in the R package *metafor*^45^. The response variable was the estimated slope of the population trend for each species, with its associated sampling variance. Separate models were fitted for each proxy of sexual selection (mating system, sexual dichromatism, sexual size dimorphism and residual testes mass). To test whether mortality modulates the effects of sexual selection, we fitted models including the interaction between each sexual selection proxy and migratory status. We predicted weaker associations in migratory species, where elevated mortality may reduce the opportunity for selection^23,27^. To test whether fecundity modifies these relationships, we fitted additional models including interactions between each sexual selection proxy and clutch size or clutch number.

All models included region (Europe and North America) as a fixed effect to account for differences in data structure and estimation. Given that there are some species that are found in both Europe and North America, we included both species and phylogeny as random effects in all models. The phylogeny was obtained from the Open Tree of Life (https://tree.opentreeoflife.org/) using the package *rotl*^46^, and branch lengths were computed using Grafen’s method with the package *ape*^47^. All analyses were performed in R version 4.5.2^48^.

## Results

The number of species with increasing and decreasing population trends was slightly biased towards declining populations; 54% of species (341 out of 738) show declining population trends. The proportion of declining populations was higher in Europe (62% species) than in North America (52% species). However, sexual selection did not affect species population trends when using either mating system (estimate ± SE) (*-0*.*0001* ± *0*.*002, p = 0*.*95, n = 658*), sexual dichromatism (*0*.*0005 ± 0*.*001, p = 0*.*77, n = 666*), sexual size dimorphism (*0*.*002 ± 0*.*006, p = 0*.*76, n = 664*), or residual testes mass (*-0*.*003 ± 0*.*002, p = 0*.*16, n = 382*) (Fig. 1). In none of the models was there an effect of region (Table S1).

**Figure 1.**
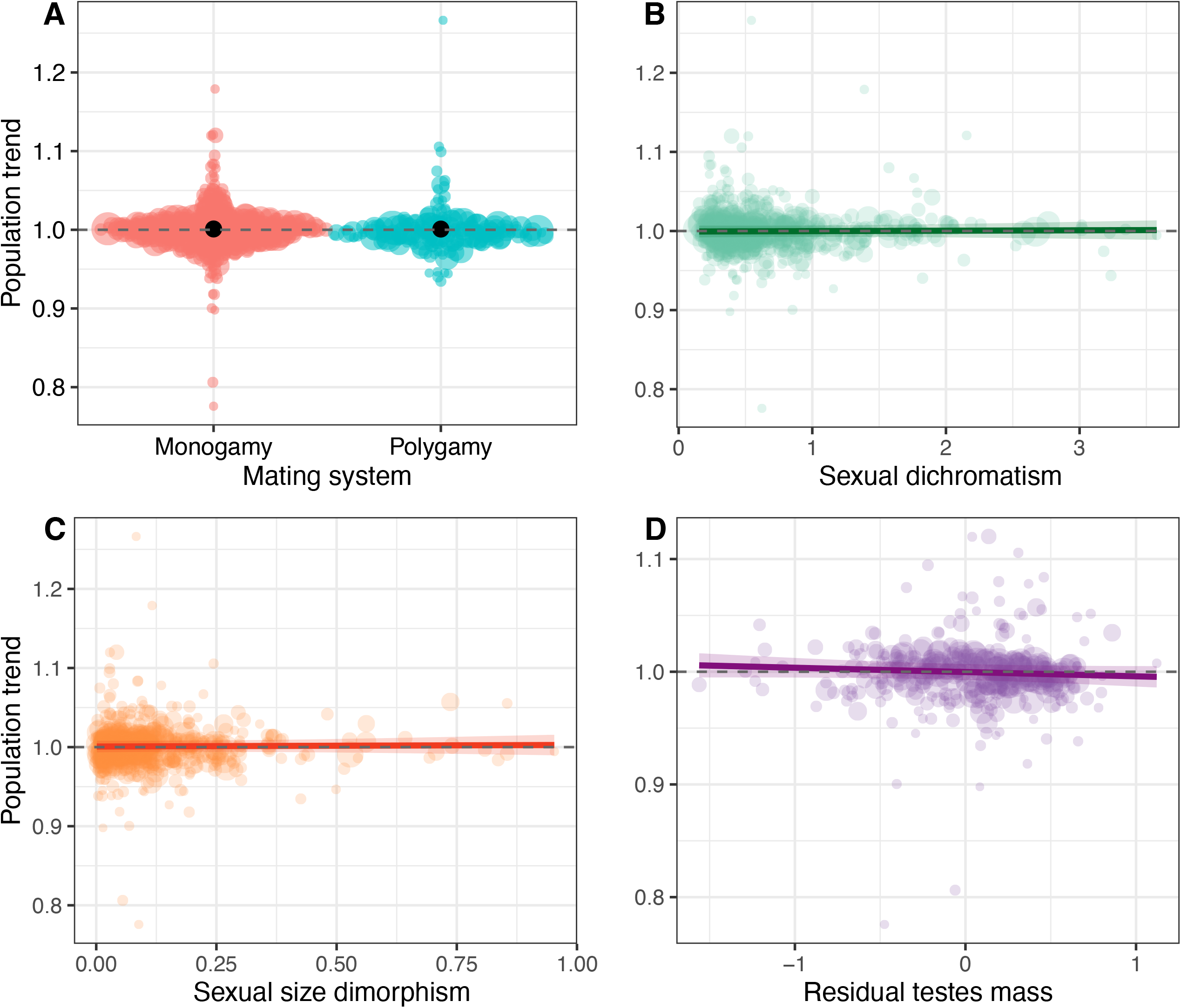
Effect of sexual selection in long-term population trends. There was no significative effect of sexual selection in long-term population trends when measured as mating system (**A**), sexual dichromatism (**B**), sexual size dimorphism, or (**C**) residual testes mass (**D**). Coloured circles represent individual species population trends with size scaled to the inverse of the variance in population trend. Black points in A and lines in B-D show the model fits and shaded are the 95% confidence interval.

When considering migratory status, we found no effect of the interaction between migratory status and sexual selection when using either mating system (-*0*.*001 ± 0*.*004, p = 0*.*72, n = 563*), dichromatism (*0*.*002 ± 0*.*003, p = 0*.*51, n = 564*), sexual size dimorphism (-*0*.*005 ± 0*.*013, p = 0*.*69, n = 578*), or residual testes mass (*0*.*001 ± 0*.*005, p = 0*.*78, n = 382*) (Fig. 2, Table S2).

**Figure 2.**
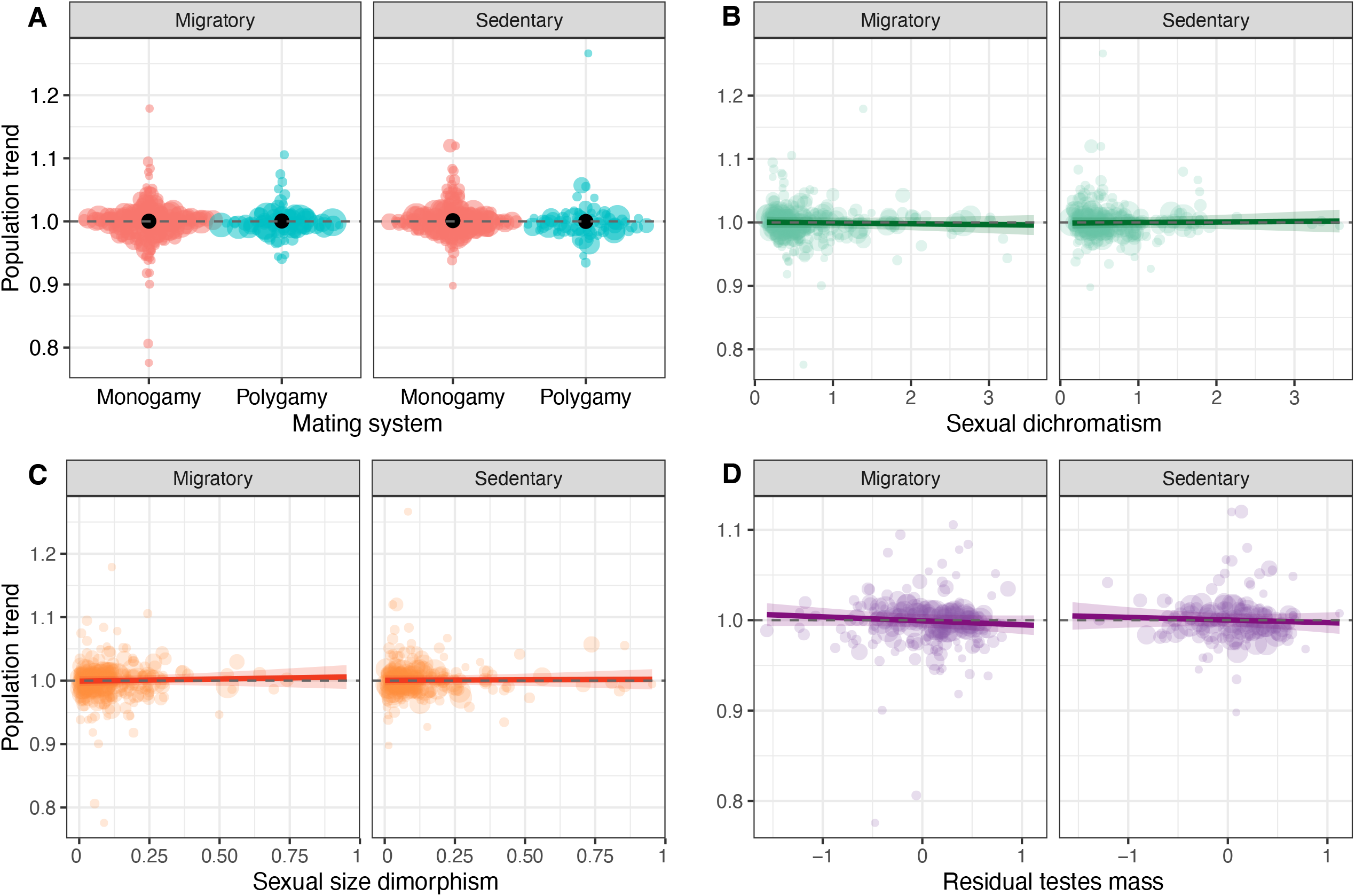
Mortality modulating the effects of sexual selection. We found no effect of sexual selection on population trends in either migratory (high offspring mortality) or resident (low offspring mortality) species when measuring mating system (**A**), sexual dichromatism (**B**), sexual size dimorphism (**C**), or residual testes mass (**D**). Coloured circles represent individual species population trends with size scaled to the inverse of the variance in population trend. Black points in A and lines in B-D show the model fits and shaded are the 95% confidence interval.

**Figure 3.**
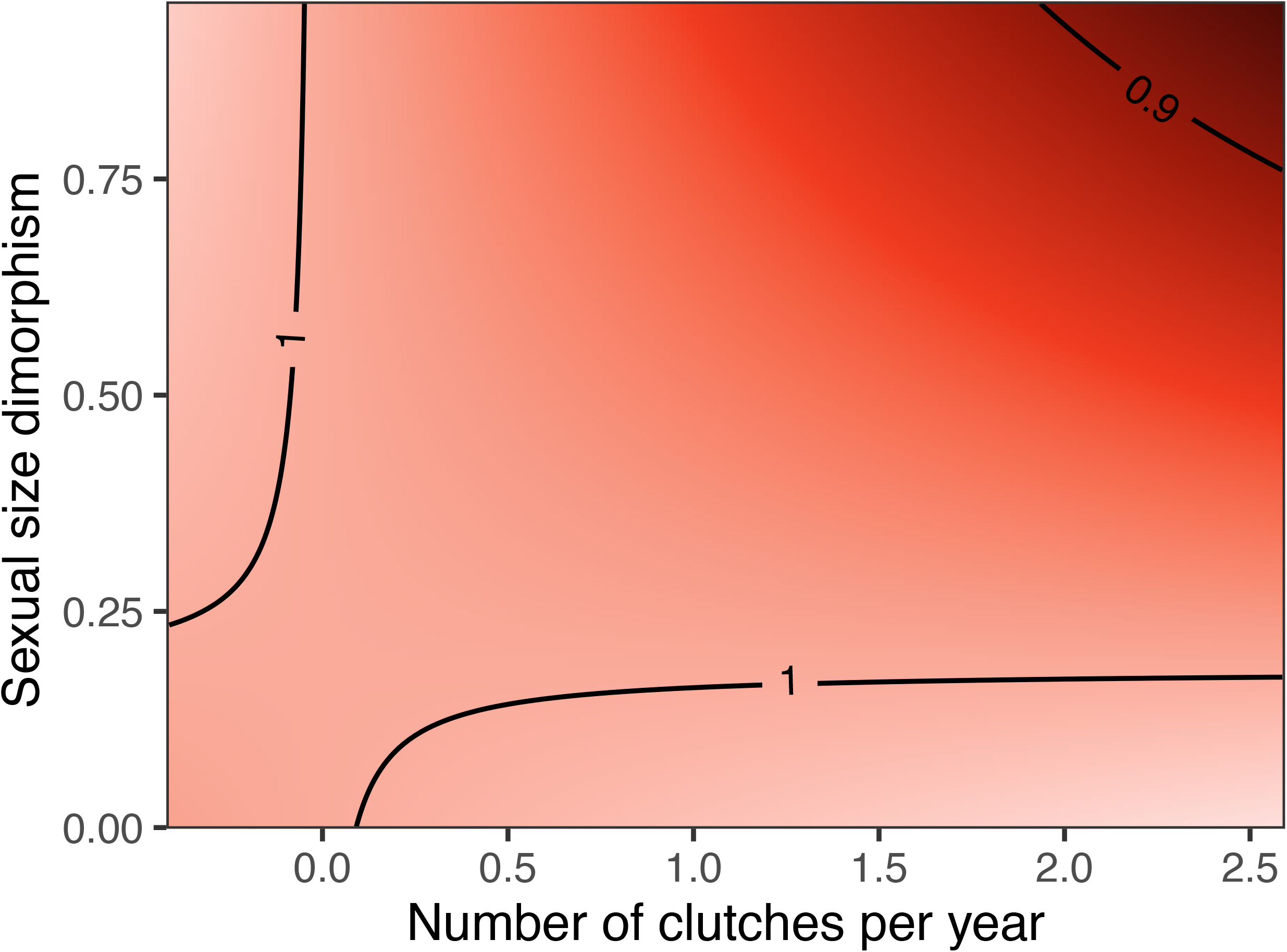
Fecundity modulates the effects of sexual selection. We found a negative effect of the interaction between clutch number and sexual size dimorphism, indicating that in highly fecund species, increased sexual selection has a more negative effect (darker colour) on long-term population trends.

We analysed if fecundity modulated the effects of sexual conflict using clutch size and clutch number. We found no significant effect of the interaction of mating system and fecundity measured either as clutch size (0.0006 ± 0.0008, *p* = 0.43, *n* = 643) or clutch number (0.0003 ± 0.003, *p* = 0.94, *n* = 593), nor the interaction of fecundity with dichromatism (clutch size: −0.0001 ± 0.0007, *p* = 0.87, *n* = 652; clutch number: 0.002 ± 0.003, *p* = 0.48, *n* = 598), or residual testes mass (clutch size: 0.0004 ± 0.001, *p* = 0.72, *n* = 381; clutch number: −0.005 ± 0.005, *p* = 0.36, *n* = 357). When using sexual size dimorphism we found no effect of the interaction with clutch size (0.003 ± 0.002, *p* = 0.22, *n* = 652) but a small and statistically significant effect of the interaction with clutch number, although in the opposite direction to that we expected (−0.032 ± 0.016, *p* = 0.043, *n* = 605) (Table S3, Table S4).

## Discussion

The role of sexual selection in shaping population fitness remains highly debated. While sexual selection can be beneficial by promoting local adaptation, its effects on population trends may be offset, or even reversed, by sexual conflict or if natural selection acts in the opposite direction^49^. Here, we analysed long-term population trends across more than 700 bird species from Europe and North America and found no evidence that sexual selection predicts long-term population trends at macroecological scales. This absence of an effect was consistent across four complementary proxies capturing both pre- and post-copulatory sexual selection, across two continents, and across contrasting demographic contexts. Together, these results suggest that, despite its central role in shaping phenotypic evolution, sexual selection does not translate into detectable effects on long-term population trajectories in natural populations.

Our results contrast with experimental studies that report small but positive effects of sexual selection on traits associated with population fitness^13^. However, they align with the majority of field-based studies, which often find null or negative relationships between sexual selection and extinction risk or population persistence^14–19^, with positive effects restricted to highly altered environments^18^. One explanation for this discrepancy is that laboratory studies typically minimize natural selection and environmental heterogeneity, which both may override the benefits of sexual selection under less benign, more natural settings^21,50–52^. Natural populations experience complex and fluctuating environments where multiple selective pressures act simultaneously and heterogeneously, potentially obscuring or counteracting the demographic consequences of sexual selection^22^. Moreover, experimental studies in laboratory settings are often biased toward species with short generation times and high fecundity, which may enhance the adaptive benefits of sexual selection compared to relatively long-lived organisms such as birds^13^.

Several non-mutually exclusive mechanisms may explain why sexual selection does not affect population trends at the large spatial and temporal scales of our study. First, the effects of sexual selection on population fitness may simply be too small relative to the magnitude of ecological and environmental variation. Population trends in the wild are driven by a combination of factors including habitat change, climate variability, anthropogenic pressures, and stochastic demographic processes. Previous research suggests that the benefits of sexual selection, if any, may only emerge under conditions of rapid or extreme environmental change, such as ongoing climate change^21^. At the continental scale of our analysis, substantial variation in the magnitude and rate of environmental change experienced among species may obscure such context-dependent effects. Future work explicitly testing how the effects of sexual selection vary with different rates of environmental change could clarify whether its benefits are restricted to more rapidly changing environments or whether, against this background of high environmental variability, the demographic consequences of sexual selection are negligible.

Second, sexual selection maximizes individual reproductive success rather than population growth per se, and these objectives are not necessarily aligned. Traits favoured by sexual selection can impose survival costs or exacerbate sexual conflict^10,11,53^, potentially offsetting any benefits arising from the spread of advantageous alleles^12,54,55^. As a result, even if sexual selection accelerates adaptation, its net effect on population growth may be neutral or highly context dependent. Third, there may be a mismatch in timescales between evolutionary and demographic processes, such that that positive effects of sexual selection via more efficient purging deleterious alleles may lag behind demographic effects induced by environmental change. This might be especially pronounced in rapidly changing environments, where the set of deleterious alleles can change faster than sexual selection can efficiently purge them.

We explicitly tested the hypothesis that mortality modulates the effects of sexual selection by comparing migratory and resident species. Increased mortality, particularly in early life stages, can reduce the opportunity for selection, potentially offsetting the benefits of sexual selection^23,27^. Migration is associated with elevated and often stochastic mortality, especially in juveniles^30^. We therefore predicted that increased mortality in migratory species would reduce any benefits of sexual selection, whereas such benefits would be more detectable in resident species. Contrary to expectations, we found no evidence that sexual selection had stronger effects in residents than in migratory species. This suggests either that mortality rates in natural populations, particularly if near their carrying capacity, are sufficient to limit the demographic consequences of sexual selection^21^, or that the effects of sexual selection are inherently weak relative to other sources of variation. Although migratory status is an indirect proxy for mortality, it captures well-established differences in survival regimes within partially migratory populations (with resident and migratory individuals)^31^, and the consistency of our results across both groups strengthens this conclusion. However, it is unclear if this difference in mortality within populations reflects patterns between species, and results should be interpreted with this caveat in mind.

Similarly, we found little support for the hypothesis that fecundity could enhance the detectability of the benefits of sexual selection. Higher fecundity is expected to buffer the demographic costs of sexual selection, thereby allowing for sexual selection to promote adaptation^18,20,23^. However, neither clutch size nor clutch number moderated the relationship between sexual selection and population trends. The only exception was a weak interaction between sexual size dimorphism and clutch number, and this effect was in the opposite direction to that predicted. Taken together, these results suggest that increased reproductive output does not translate into stronger demographic effects of sexual selection, at least in birds.

Our study has limitations that should be considered when interpreting these results. First, although we used multiple proxies of sexual selection, including mating system, sexual dichromatism, sexual size dimorphism, and residual testes mass, all proxies are imperfect and may capture aspects of natural selection in addition to sexual selection^24,26^. However, the consistency of our results across proxies suggests that this limitation is unlikely to explain the absence of an effect. Second, although we harmonized population trend estimates between European and North American datasets using comparable modelling approaches, residual differences in sampling design and data structure may introduce additional noise. Nevertheless, we found no effect of region in any model, suggesting that methodological differences did not bias our conclusions.

Overall, our results highlight a potential disconnect between evolutionary processes and population-level dynamics. Sexual selection is a powerful driver of phenotypic diversification and adaptation^1^, yet its effects on population trajectories appear to be negligible or highly context-dependent at large spatial and temporal scales. Ultimately, our results suggest that sexual selection might not promote the persistence of species, which is in line with findings that sexual selection is typically only weakly associated with species diversity in birds^56^ and other taxonomic groups^57^. One possibility is that the benefits of sexual selection only emerge under specific conditions, such as rapid environmental change or population bottlenecks, where adaptation is strongly limiting population persistence. At the continental scale considered here, variation in environmental conditions and demographic processes may obscure such context-dependent effects. Future research integrating fine-scale demographic data, experimental approaches, and explicit measures of environmental change will be essential to determine under which conditions sexual selection can affect population trends.

## Acknowledgements

EHM was supported by the Swedish Research Council (grant number: 2025-03964). CRC was supported by a Natural Environment Research Council Independent Research Fellowship (NE/T01105X/1). TJ was supported by the French National Research Agency (ANR-25-CE02-1229).

