## Supplementary Information for "Sexual selection does not predict long-term population trends in birds"

**Table S1.** Sexual selection had no effect in long-term population trends across birds either when measured as mating system (A), sexual dischromatism (B), residual testes mass (C) nor sexual size dimorphism (D).

| | Estimate (SE) | t value | df | p |
|---|---|---|---|---|
| **A) Mating system** | | | | |
| Intercept (Monogamy) | 0.996 (0.005) | 173.48 | 599 | < 0.001 *** |
| Region (North America) | 0.004 (0.004) | 0.94 | 655 | 0.34 |
| Polygamy | -0.0001 (0.002) | -0.05 | 655 | 0.95 |
| **B) Sexual dichromatism** | | | | |
| Intercept | 0.995 (0.005) | 185.1 | 609 | < 0.001 *** |
| Region (North America) | 0.003 (0.003) | 0.95 | 663 | 0.33 |
| Sex dichromatism | 0.0005 (0.001) | 0.28 | 609 | 0.77 |
| **C) Sexual size dimorphism** | | | | |
| Intercept | 0.996 (0.005) | 185.87 | 601 | < 0.001 *** |
| Region (North America) | 0.004 (0.004) | 1.05 | 661 | 0.29 |
| Sexual size dimorphism | 0.002 (0.006) | 0.29 | 601 | 0.76 |
| **D) Residual testes mass** | | | | |
| Intercept | 0.997 (0.006) | 162.62 | 335 | < 0.001 *** |
| Region (North America) | 0.002 (0.005) | 0.38 | 379 | 0.7 |
| Redisual testes mass | -0.003 (0.002) | -1.39 | 335 | 0.16 |

**Table S2.** Mortality, measured as migratory status, did not show an effect mediating the effects of sexual selection in long-term population trends.

| | Estimate (SE) | t value | df | p |
|---|---|---|---|---|
| **A) Mating system** | | | | |
| Intercept | 0.996 (0.006) | 165.001 | 510 | < 0.001 *** |
| Region (North America) | 0.003 (0.004) | 0.82 | 558 | 0.41 |
| Polygamy | 0.0003 (0.003) | 0.09 | 558 | 0.92 |
| Migratory status (Sedentary) | 0.0008 (0.002) | 0.404 | 510 | 0.68 |
| Polygamy : Sedentary | -0.001 (0.004) | -0.35 | 558 | 0.72 |
| **B) Sexual dichromatism** | | | | |
| Intercept | 0.99 (0.006) | 512 | 165.86 | < 0.001 *** |
| Region (North America) | 0.003 (0.004) | 559 | 0.87 | 0.38 |
| Sex dichromatism | -0.001 (0.002) | 512 | -0.54 | 0.58 |
| Migratory status (Sedentary) | -0.001 (0.003) | 512 | -0.52 | 0.60 |
| Dichromatism : Sedentary | 0.002 (0.003) | 512 | 0.64 | 0.51 |
| **C) Sexual size dimorphism** | | | | |
| Intercept (Monogamy) | 0.995 (0.005) | 176.00 | 520 | < 0.001 *** |
| Region (North America) | 0.003 (0.004) | 0.84 | 573 | 0.40 |
| Sexual size dimorphism | 0.006 (0.01) | 0.65 | 520 | 0.51 |
| Migratory status (Sedentary) | 0.001 (0.002) | 0.55 | 520 | 0.57 |
| Dimorphism: Sedentary | -0.005 (0.013) | -0.39 | 520 | 0.69 |
| **D) Residual testes mass** | | | | |
| Intercept (Monogamy) | 0.997 (0.006) | 159.49 | 333 | < 0.001 *** |
| Region (North America) | 0.002 (0.005) | 0.39 | 377 | 0.69 |
| Redisual testes mass | -0.004 (0.003) | -1.27 | 333 | 0.203 |
| Migratory status (Sedentary) | 0.0009 (0.002) | 0.39 | 333 | 0.69 |
| Testes mass : Sedentary | 0.001 (0.005) | 0.27 | 333 | 0.78 |

**Table S3.** Fecundity, measured as clutch size, had no effect mediating the effects of sexual selection in long-term population trends

| | Estimate (SE) | t value | df | p |
|---|---|---|---|---|
| **A) Mating system** | | | | |
| Intercept (Monogamy) | 0.997 (0.005) | 176.19 | 582 | < 0.001 *** |
| Region (North America) | 0.003 (0.004) | 0.81 | 638 | 0.41 |
| Clutch size | -0.001 (0.0006) | -1.82 | 582 | 0.068 |
| Polygamy | -0.001 (0.001) | -0.58 | 638 | 0.56 |
| Clutch size : Polygamy | 0.0006 (0.0008) | 0.78 | 638 | 0.43 |
| **B) Sexual dichromatism** | | | | |

| | | | | |
|---|---|---|---|---|
| Intercept | 0.996 (0.005) | 185.21 | 593 | < 0.001 *** |
| Region (North America) | 0.003 (0.003) | 0.82 | 647 | 0.4 |
| Clutch size | -0.0008 (0.0007) | -1.07 | 593 | 0.28 |
| Sex dichromatism | 0.0005 (0.001) | 0.24 | 593 | 0.8 |
| Clutch size : Dichromatism | -0.0001 (0.0007) | -0.15 | 593 | 0.87 |
| **C) Sexual size dimorphism** | | | | |
| Intercept | 0.996 (0.005) | 185.58 | 587 | < 0.001 *** |
| Region (North America) | 0.003 (0.004) | 0.87 | 647 | 0.38 |
| Clutch size | -0.001 (0.0006) | -2.09 | 587 | 0.03 |
| Sexual size dimorphism | 0.001 (0.006) | 0.28 | 587 | 0.77 |
| Clutch size : Dimorphism | 0.003 (0.002) | 1.22 | 587 | 0.22 |
| **D) Residual testes mass** | | | | |
| Intercept | 0.998 (0.006) | 158.21 | 332 | < 0.001 *** |
| Region (North America) | 0.001 (0.005) | 0.3 | 376 | 0.75 |
| Clutch size | -0.001 (0.0007) | -1.43 | 332 | 0.15 |
| Redisual testes mass | -0.003 (0.002) | -1.37 | 332 | 0.17 |
| Clutch size : Testes mass | 0.0004 (0.001) | 0.35 | 332 | 0.72 |

**Table S4.** Fecundity, measured as clutch number, had no effect mediating the effects of sexual selection in long-term population trends, except for a negative interaction with sexual size dimorphism.

| | **Estimate (SE)** | **t value** | **df** | **p** |
|---|---|---|---|---|
| **A) Mating system** | | | | |
| Intercept (Monogamy) | 0.996 (0.005) | 182.93 | 533 | < 0.001 *** |
| Region (North America) | 0.003 (0.004) | 0.8 | 588 | 0.41 |
| Clutch number | 0.003 (0.002) | 1.74 | 533 | 0.082 |
| Polygamy | -0.002 (0.002) | -1.05 | 588 | 0.29 |
| Clutch No : Polygamy | 0.0003 (0.003) | 0.07 | 588 | 0.94 |
| **B) Sexual dichromatism** | | | | |
| Intercept | 0.995 (0.005) | 194.44 | 540 | < 0.001 *** |
| Region (North America) | 0.004 (0.004) | 1,00 | 593 | 0.31 |
| Clutch number | 0.002 (0.002) | 0.7 | 540 | 0.47 |
| Sex dichromatism | 0.0001 (0.001) | 0.03 | 540 | 0.97 |
| Clutch No : Dichromatism | 0.002 (0.003) | 0.7 | 540 | 0.48 |
| **C) Sexual size dimorphism** | | | | |
| Intercept | 0.994 (0.004) | 205.82 | 541 | < 0.001 *** |
| Region (North America) | 0.004 (0.004) | 1.12 | 600 | 0.25 |
| Clutch number | 0.005 (0.002) | 2.71 | 541 | 0.006 ** |

| | | | | |
|---|---|---|---|---|
| Sexual size dimorphism | -0.0007 (0.007) | -0.10 | 541 | 0.91 |
| Clutch No : Dimorphism | -0.032 (0.016) | -2.02 | 541 | 0.043 * |
| **D) Residual testes mass** | | | | |
| Intercept | 0.997 (0.005) | 170.44 | 309 | < 0.001 *** |
| Region (North America) | 0.002 (0.005) | 0.37 | 352 | 0.7 |
| Clutch number | 0.006 (0.002) | 2.71 | 309 | 0.007 ** |
| Redisual testes mass | -0.005 (0.003) | -1.87 | 309 | 0.061 |
| Clutch No : Testes mass | -0.005 (0.005) | -0.9 | 309 | 0.36 |
